# ChemGAPP; A Package for Chemical Genomics Analysis and Phenotypic Profiling

**DOI:** 10.1101/2023.01.05.522861

**Authors:** Hannah M. Doherty, George Kritikos, Marco Galardini, Manuel Banzhaf, Danesh Moradigaravand

**Affiliations:** University of Birmingham, Institute of Microbiology and Infection and School of Biosciences, B15 2TT Birmingham, UK; Genome Biology Unit, European Molecular Biology Laboratory, Heidelberg, Germany; Institute for Molecular Bacteriology, TWINCORE Centre for Experimental and Clinical Infection Research, a joint venture between the Hannover Medical School (MHH) and the Helmholtz Centre for Infection Research (HZI), Hannover, Germany; Cluster of Excellence RESIST (EXC 2155), Hannover Medical School (MHH), Hannover, Germany; Biological and Environmental Science and Engineering (BESE) Division, King Abdullah University of Science and Technology (KAUST), Saudi Arabia

**Keywords:** Chemical Genomics, Bacteria, Genetics, Quality Control Analysis, Automated Data Analysis

## Abstract

High-throughput chemical genomic screens produce informative datasets, providing valuable insights into unknown gene function on a genome-wide level. However, there is currently no comprehensive analytic package publicly available. We developed and benchmarked ChemGAPP to bridge this gap. ChemGAPP allows integration of various steps in a streamlined and user-friendly format, including rigorous quality control measures to curate screening data. ChemGAPP provides three sub-packages for different chemical-genomic screens: ChemGAPP Big for handling large-scale high-throughput screens; ChemGAPP Small, designed for small-scale screen analysis and ChemGAPP GI for genetic interaction screen analysis. ChemGAPP is available at https://github.com/HannahMDoherty/ChemGAPP.

## Background

The field of chemical genomics and high throughput phenomic profiling has revolutionised the ability to functionally annotate unknown genes on a genome wide level. With applications for drug discovery, mechanism of action studies and conditional essentiality studies, chemical genomics is a rapidly growing field ^1,2^. Chemical genomic screens systematically assess the effect of chemical or environmental perturbations (stresses) on single-gene mutant libraries. The resulting phenotypes can range from colony size as a proxy of fitness, colour uptake to quantify biofilm formation, and changes in colony topology to asses biofilm morphology ^3^. Individual observations can be studied in isolation, based on the observed phenotype of a gene in a specific condition, thereby creating a functional link between a stress and a defined genetic perturbation. However, the true power of chemical genomic approaches lays in the ability to calculate phenotypic profiles for each mutant based on their phenotypes across conditions ^4^. These phenotypic profiles can be hierarchically clustered to reveal similarities between mutant profiles to functionally cluster genes or stress conditions. This linkage reconstitutes biological pathways and complexes and therefore can functionally cluster unknown genes to known biology ^4^. The power of chemical genomics screens, and the plethora of valuable information they produce, has led to many important biological discoveries and their popularity in the field of systems microbiology ^4–12^. The first major chemical genomic screen within bacteria was performed in *Escherichia coli*. Nichols et al., in 2011, assessed the phenotypes of the KEIO collection, an in-frame single-gene knockout mutant library in *E. coli* K-12, within over 300 conditions ^4,13^. The Nichols et al., study successfully produced >10,000 phenotypes and categorised various biologically relevant hits and suggested functions for genes with previously unknown function ^4^.

Despite the rise in chemical genomics studies, there are currently no dedicated programs to analyse the type of data these screens produce. Currently, a variety of methods are being implemented for the analysis of data and are being performed by in-house scripts or are adaptations of packages used for similar techniques ^4,14–18^. A number of image analysis software are available for the reading of chemical genomic screening plates, such as gitter or Iris ^3,19^. Gitter and Iris convert phenotypes within plate images into numerical values, however they are unable to convert these values into fitness scores ^3,19^. Another software that has been frequently implemented, which is able to produce fitness scores is EMAP Toolbox, however, this tool is now deprecated ^4,17,20^. Furthermore, EMAP was originally developed to handle genetic interactions within yeast and henceforth does not offer a standard solution for handling data from assays under conditions. None of these options are accessible for users with limited computational expertise, as many require the knowledge of coding languages such as R or MATLAB to implement ^15,16,18^. Consequently, there is a clear gap that hinders researchers from effectively using chemical genomics approaches. This study, therefore, introduces an easy to use, new wrapper software and streamlit app called ChemGAPP (Chemical Genomics Analysis and Phenotypic Profiling), which has been developed as a comprehensive chemical genomics data analysis software.

Here we provide evidence for ChemGAPP’s effectiveness to reliably analyse chemical genomics data. By providing ChemGAPP it will enable the wider scientific community to use chemical genomics approaches to reveal significant, biologically relevant insights into gene function, drug mechanisms of action and antibiotic resistance mechanisms.

## Results

### ChemGAPP Pipelines

ChemGAPP comprises of three pipelines ChemGAPP Big, ChemGAPP Small and ChemGAPP GI, each specifically designed for different types of chemical genomic screens (Fig. 1).

**Figure 1:**
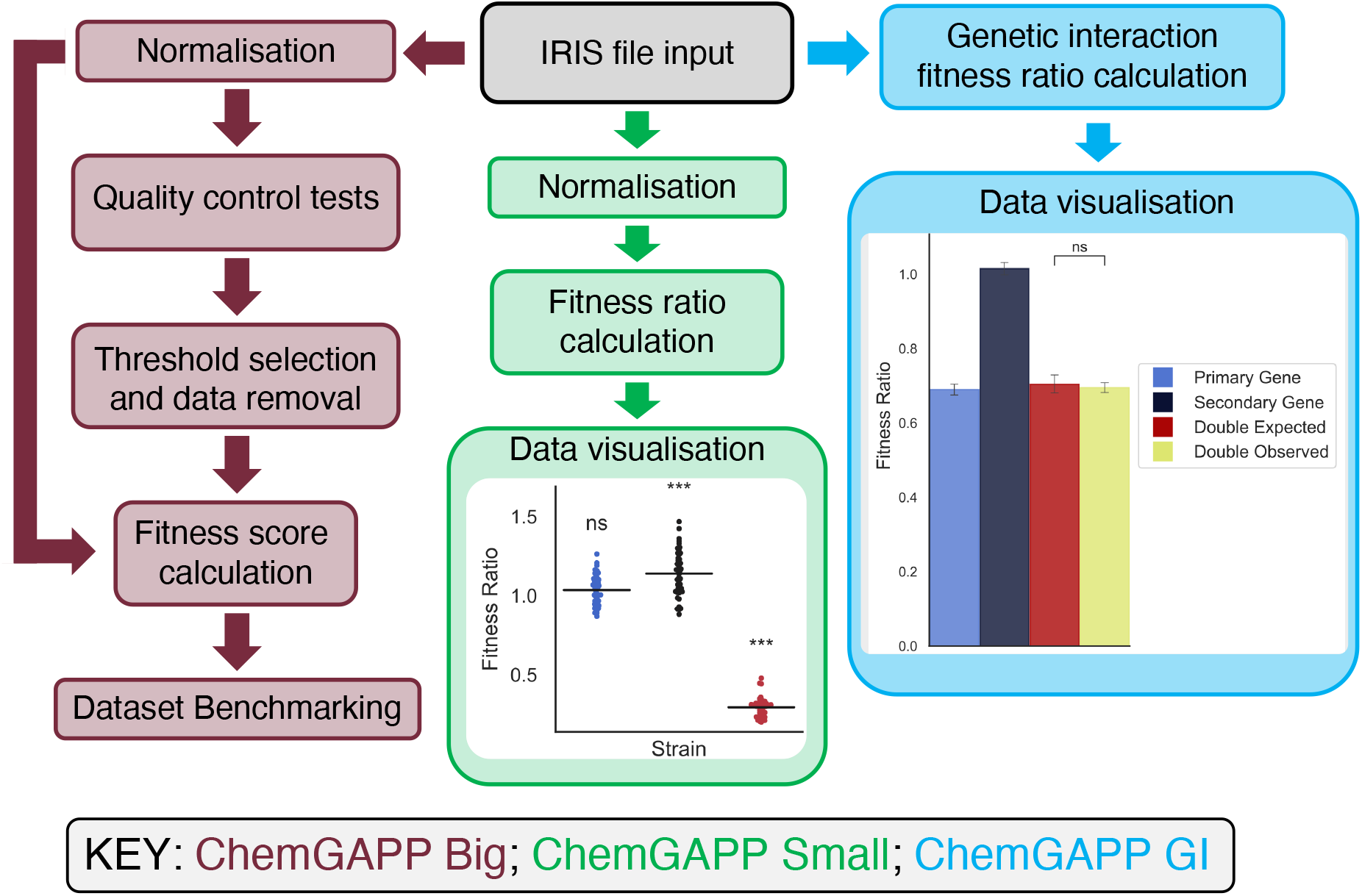
Workflow of the ChemGAPP packages. ChemGAPP Big analyses large-scale chemical genomic screens. ChemGAPP Small analyses small-scale chemical genomic screens. ChemGAPP GI analyses small-scale genetic interaction screens.

### ChemGAPP Big

ChemGAPP Big processes all classical stages of data analysis within large-scale chemical genomic screens, whilst introducing new features to improve upon this analysis. The first stage is to produce a dataset from the experimental data. The reading of chemical genomic screen data is increasingly being performed by the imaging analysis software Iris ^3^. Iris’s increasing popularity is due to its versatility in phenotype measurements. Iris can analyse many phenotypes, including colony size, integral opacity, circularity, colour, etc. ^3^. Therefore, the Iris file format was chosen as the default input for ChemGAPP Big. Despite this, there is no limit to the usage of other image analysis software, such as gitter, as long as the data is compiled into the Iris file format ^19^.

ChemGAPP Big then removes noise from the experimental data and scales data to become comparable via a two-step normalisation. It is vital to perform plate normalisation before the data can be scored, due to the noise that arises during large chemical screens. A major issue within chemical genomic screens in plate format is the edge effect, which is targeted by the first stage of normalisation ^21^. Since mutants are densely pinned on plates, the outermost colonies are left with more space. This results in increased growth around the plate edges, due to reduced nutrient competition ^21^. The second stage standardises all plates such that regardless of condition, the median colony sizes are comparable. This is important for the generation of mutant fitness scores across different conditions later in the pipeline.

Within ChemGAPP Big, we have implemented multiple quality control tests to find common errors which occur within chemical genomic screens. Acquiring the physical raw data for these screens is challenging as often thousands of plates need to be accurately processed. The most common errors which occur in the laboratory are mislabelled plates, inverted images, and mutants being unequally pinned between replicates or missed entirely. In addition, despite Iris’s ability to robustly quantify most mutants, its detection algorithm can fail and create artefacts. The combination of the quality control tests is flexible, giving the user control over the curation of the dataset. Providing the user with the ability to employ the right tests for the issues that have arisen within their datasets. Curation of the dataset is a fine balance between removing detrimental data, but not removing so much data that the dataset is no longer informative. Therefore, in order to provide flexibility, the thresholds at which replicate plates and conditions are removed is left to the user. The user can equally choose to skip curation if the quality control tests reveal a clean dataset. However, ChemGAPP Big allows for an informed choice. ChemGAPP Big outputs a visual representation of the quantity of data that will be lost at different thresholds for the various tests (Fig. S1).

In order to find meaning within the data, fitness scores must be produced. Within ChemGAPP Big this is performed using the S-score test. The S-score test, as described in Collins et al., 2006, is a modified T-test where the mean of a set of colony replicates, for the desired mutant within a specific condition, are compared to the median size of that mutant across all conditions ^17^. Thus, providing a statistical score for fitness, with positive scores showing increased fitness and negative scores showing decreased fitness. Comparing a mutant to itself across all conditions as a simulated wildtype is more powerful than comparing to the true wildtype. This is because a mutant may have generally increased or decreased fitness than the wildtype. By comparing to itself, a true baseline for comparison is used. Furthermore, since S-scores correlate to statistical significance, they provide a robust and informative measurement of how strong a phenotype is across various conditions. Both the curated and non-curated datasets are then scored by ChemGAPP Big.

Following dataset curation, it is vital to confirm which dataset has the most accurate fitness scores. In order to measure the accuracy of the datasets, the concept that genes within the same operon will often have a similar function was drawn upon. If genes are functionally related, we would expect their knockout mutants to behave similarly across the different tested conditions. Thus, they would have similar phenotypic profiles, which is their set of S-scores across all conditions ^4^. ChemGAPP Big compares genes from the same operon and different operons and produces a receiver operating characteristic (ROC) curve for both the curated and non-curated datasets for comparison. The dataset with higher similarity between genes from the same operon can then be determined as the more predictive dataset and should be used as the final dataset for the study.

### ChemGAPP Small

In large-scale chemical genomics, by design, the screens are unbiased, and the aim is to test mutant libraries against a manifold of stresses. Since in most conditions mutants are unlikely to display a phenotype, a normal distribution of colony sizes is achieved. By contrast in small-scale studies a few or only one condition are screened, and, in such analyses, a confined hypothesis is often being tested. When testing a specific hypothesis, conditions are often specifically chosen with the assumption that there will be an associated phenotype. Therefore, the conjecture about the normal distribution of fitness effects across conditions may not hold. In these instances, ChemGAPP Big is not fit for purpose.

Therefore, we developed ChemGAPP small for the analysis of targeted small-scale chemical genomics studies. ChemGAPP Small generates fitness ratios for mutants versus the wildtype on the corresponding condition, in place of S-scores, without relying on any assumption about fitness distributions across conditions. ChemGAPP Small then calculates the statistical significance of the fitness ratios using a one-way ANOVA. By doing so, ChemGAPP Small allows users to robustly analyse smaller datasets without any constraints on gene or condition number.

### ChemGAPP GI

The final package ChemGAPP GI (ChemGAPP <u>G</u>enetic <u>I</u>nteractions) can analyse genetic (epistatic) interaction screening data. ChemGAPP GI is able to analyse single or multiple gene pairs on the same condition plate, producing separate plots for each gene pair. Genetic interaction studies aim to determine the type of epistasis between two genes. Genetic interactions are defined as instances where the observed fitness of a double knockout mutant is significantly different to the expected fitness ratio ^22^. This can be further categorised into two types, positive (alleviating), and negative (synergistic). In positive epistasis the double knockout is fitter than anticipated and in negative epistasis it shows decreased fitness ^22^.

ChemGAPP GI assesses the fitness of two single knockout mutants and a double knockout mutant, versus a wildtype, to calculate the observed and expected fitness ratios. By doing so, the tool not only provides a proxy for the strength of the genetic interactions but determines the mode of epistasis between the two query genes.

### ChemGAPP Big normalisation produces more uniform data for scoring

In order to assess the ChemGAPP Big pipeline, we used the chemical genomic screen data from Nichols et al., 2011, which was re-analysed using the image analysis software Iris by Kritikos et al., 2017 ^3,4^. Kritikos et al, measured colony integral opacity as the measure for colony size, since this provides a better measure of 3-dimensional colony growth ^3^. The colony integral opacity data from Nichols et al., was provided by Kritikos et al, for the current study ^3^.

To validate the effectiveness of normalisation, the Nichols et al. data was fed into ChemGAPP Big, and normalisation was performed. To evaluate if the normalisation steps reduced variation due to condition associated growth effects, the distributions of three randomly selected plates were analysed pre and post normalisation, see Figure 2. Prior to normalisation, the distributions of colony sizes were significantly different to each other, based on a Mann-Whitney test. However, upon the normalisation, the distribution of sizes became more uniform. Furthermore, the significance in their difference was lost, without losing the nuances of the data, Figure 2.

**Figure 2:**
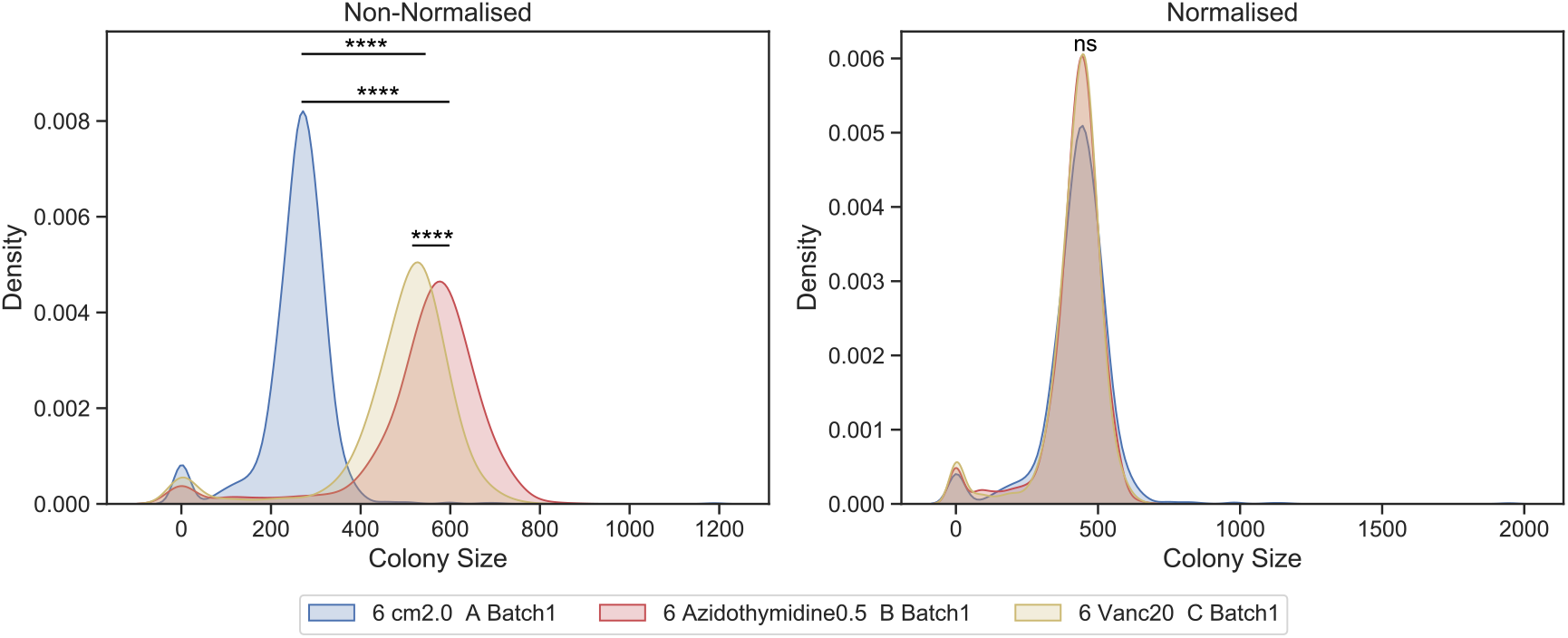
Density plot showing the difference between pre and post normalised colony sizes for three randomly selected condition plates. Cm2.0; Chloramphenicol 2 μg/mL, Azidothymidine0.5; Azidothymidine 0.5 ng/mL, Vanc20; Vancomycin 20 μg/mL. ****: p-value ≤ 0.0001, ns: p-value > 0.05.

### ChemGAPP Big identifies common errors within chemical genomic screens

In order to tailor ChemGAPP Big to large-scale chemical genomic screen data, we designed the quality control steps to target common errors that arise within the field. One frequent error is the introduction of plate effects due to unequal pinning. To identify this, we conducted the Z-score test and looked at plates with low percentage normality. The Z-score test highlights outlier colony sizes between replicate colonies. ChemGAPP Big considers non-outlier colonies as ‘normal’, therefore a plate with multiple outlier colony sizes will have a low normality percentage. Two replicate plates C and D in the 20 °C cold shock condition, had a reduced normality percentage of 64.84% and 70.53%, respectively (Fig. 3A). Figure 3A represents the colony sizes of the replicate plates for the 20 °C condition. Conversely to replicates C and D, plates A and B had high percentage normality scores of 90.43% and 95.05%, respectively (Fig. 3A). Unequal pinning can be clearly observed within plate C and D. Within D, the upper segment of the plates has consistently smaller colonies and the lower segment has consistently larger colonies. Whereas within replicate C, it is the outer segments of the plates, excluding the corners, with reduced colony sizes, and the inner segment with increased sizes. Within plate C and D, 17.9% and 15.69% of colonies were classified as larger than the mean replicate colony size, and 17.19% and 13.74% of colonies were smaller, respectively, reflecting what is seen in Figure 3A.

**Figure 3:**
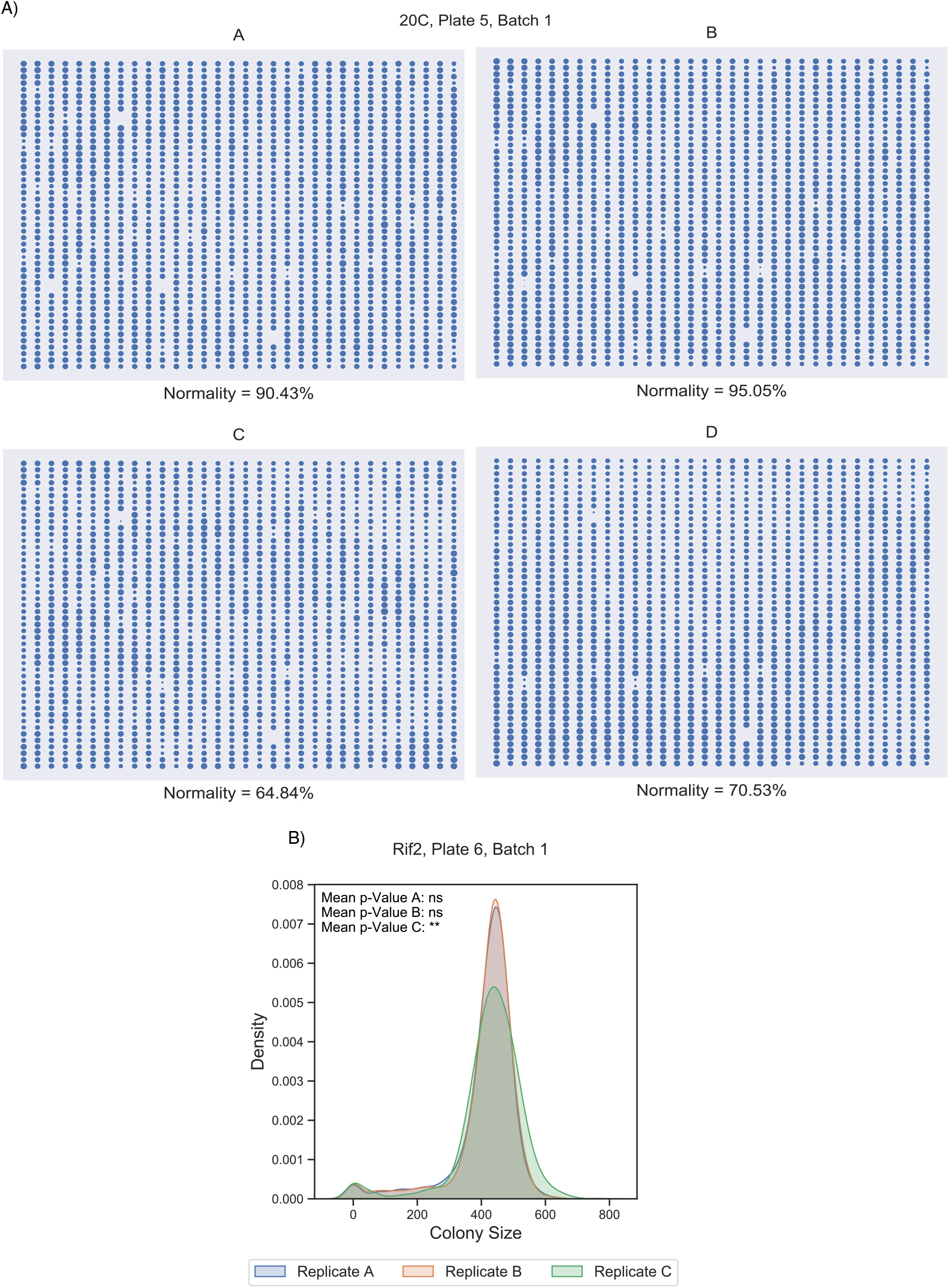
ChemGAPP Big highlights plates with errors common to chemical genomic screens. A) Plate matrix depicting the colony sizes within replicate plates of the condition 20C (cold shock 20 °C) and the percentage normality determined by the Z-score test; B) Density of colony sizes for replicates A, B and C for condition Rif2 (Rifampicin 2 μg/mL), Plate 6, Batch1. Difference in the distribution of C vs A and B is statistically significant by Mann-Whitney test. ** = 0.001 < p-value ≤ 0.01; ns = non-significant.

To further detect unequal pinning, as well as missed pinning or undetected colonies, the Mann-Whitney Test was performed. These effects are evident in Figure 3B, where we identified a condition, i.e., 2 μg/mL rifampicin, in which one replicate (C) differed in its distribution of colony sizes. Replicate C had a mean Mann-Whitney p-value of 0.0018 (Fig. 3B). The distribution of C shown in Figure 3B shows an increased number of larger and smaller colonies than A or B. Furthermore, C had more missing colonies (21) than A or B which both had only 4 missing values (Fig. S2). This is likely due to colonies being missed by pinning or by Iris detection, showing the ability of the tool to successfully pinpoint non-reproducible replicate plates based on a variety of defects.

### S-scores are reliable and accurate representations of fitness

In order to evaluate if the S-scores produced by ChemGAPP Big were accurate and robust representations of fitness, a bootstrapped dataset was produced. The non-curated and curated normalised datasets were bootstrapped and subjected to S-score analysis by ChemGAPP Big. The mean absolute error (MAE) was calculated between the mean of the bootstrapped S-scores and the non-bootstrapped S-scores for each individual score. The MAE for the non-curated dataset was 0.0622, therefore, on average the mean s-scores from the bootstrapped data differed by only 0.0622 when compared to the corresponding experimental S-score (Fig. 4A). The standard deviation (std) for the S-scores within the non-curated dataset was 1.257. The MAE constitutes just 4.94% of the error, making the MAE a negligible difference. For the bootstrapped curated dataset, the MAE was 0.0657 and the std for the original curated dataset was 1.256 (Fig. 4B). The MAE equals just 5.23% of the std, again making the difference negligible, therefore demonstrating the robustness of the S-scores.

**Figure 4:**
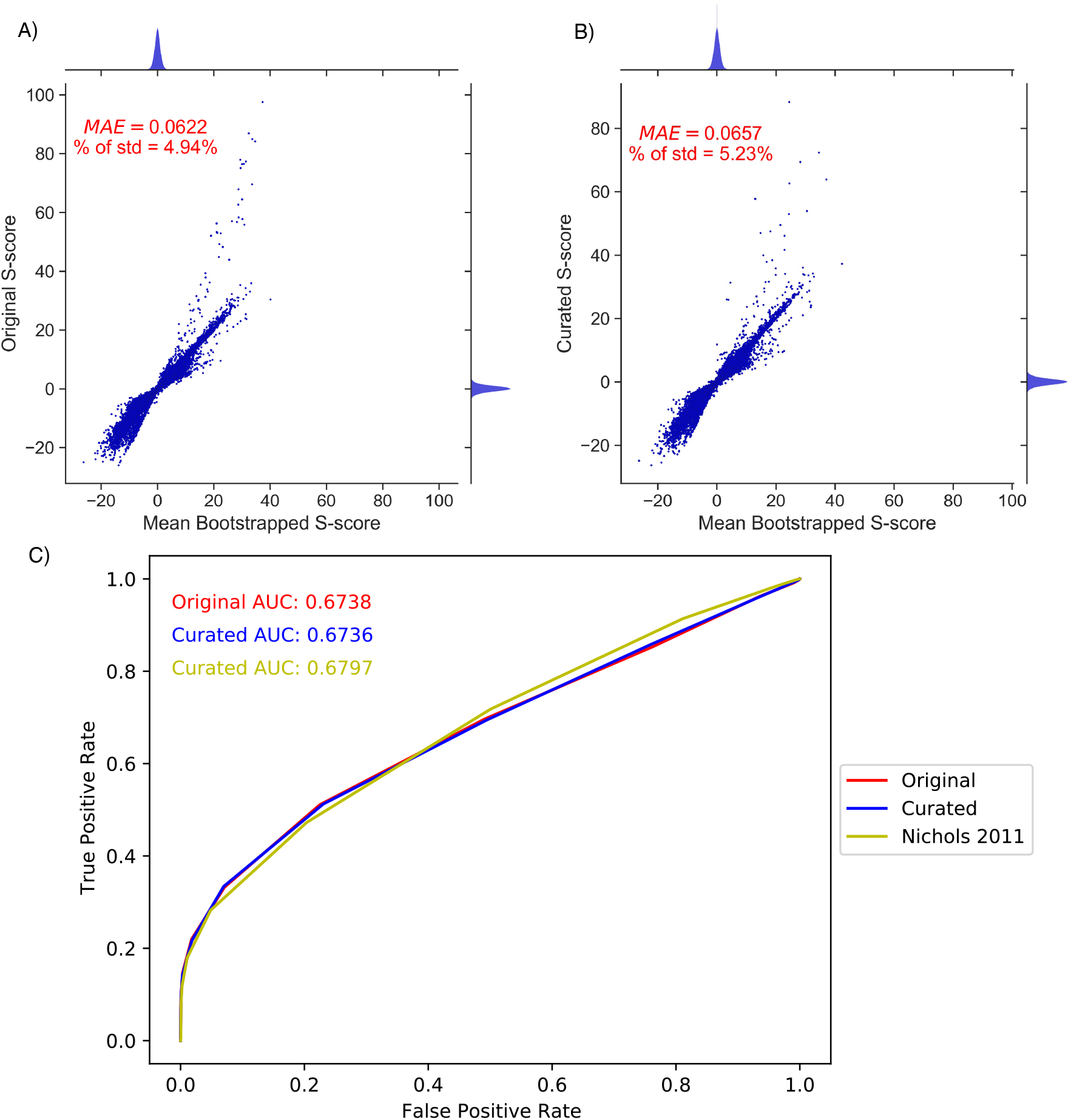
ChemGAPP produces robust and accurate S-scores. A-B) Joint scatter and density plot depicting the difference between A) the original non-curated S-scores versus the mean bootstrapped S-scores for each mutant within each condition of the KEIO dataset. B) The curated S-scores versus the mean bootstrapped S-scores for each mutant within each condition of the KEIO curated dataset. Density plots show distribution of hits with various S-scores. For ease of visualisation, outlier bootstrapped S-scores > 125 were excluded, representing a negligible 0.000018% (A), and 0.000026% (B) of all values (see Additional File 2). Outlier curated S-scores > 125 were also excluded, representing a negligible percentage of all values (0.00056%). MAE = Mean Absolute Error; % of std = Percentage of the standard deviation for the original dataset that the MAE constitutes. C) ROC curve with AUC values for the KEIO ChemGAPP Big non-curated dataset (red), the KEIO ChemGAPP Big curated dataset (blue) and the KEIO dataset from Nichols et al., 2011 (yellow).

Both the non-curated and curated datasets produced robust and reliable S-scores. However, following dataset curation, it is imperative to benchmark the fitness scoring of the curated dataset against the non-curated dataset. Furthermore, it is important to confirm ChemGAPP Big is as effective as previous software. In order to do this, a cosine similarity analysis of phenotypic profiles was performed between genes from the same operons and those from different operons. ChemGAPP Big was equally proficient at fitness score assignment as the methods employed by Nichols et al, for both the curated and non-curated datasets (Fig. 4C). The area under the curve (AUC) for the curated and non-curated datasets were 0.6738 and 0.6736, respectively, versus an AUC of 0.6797 for Nichols et al. Since some genes in the same operon may not have the same function or directly opposite functions, this measure is not a perfect model. Therefore, to further prove the effectiveness of ChemGAPP Big, it was vital to explore if biologically relevant phenotypes were still displayed within the dataset.

### ChemGAPP Big displays hits with biological significance

The major aim of ChemGAPP Big is to produce datasets which are able to accurately reveal biologically relevant phenotypes. Therefore, it is crucial that when reanalysing the Nichols et al., dataset with ChemGAPP Big that the biologically significant hits were retained. Within the Nichols et al., paper, they discovered that mutants within the GCV system were susceptible to sulfonamides ^4^. The GCV system is one of two branches responsible for the conversion of tetrahydrofolate (THF) to 5,10-methylene THF, an essential process within the tetrahydrofolate biosynthesis pathway. Nichols et al., performed validation experiments and determined this to be a true phenotype in the dataset. This phenotype was retained within the ChemGAPP Big dataset, with significant S-scores between −3 and −9 (Fig. 5A).

**Figure 5:**
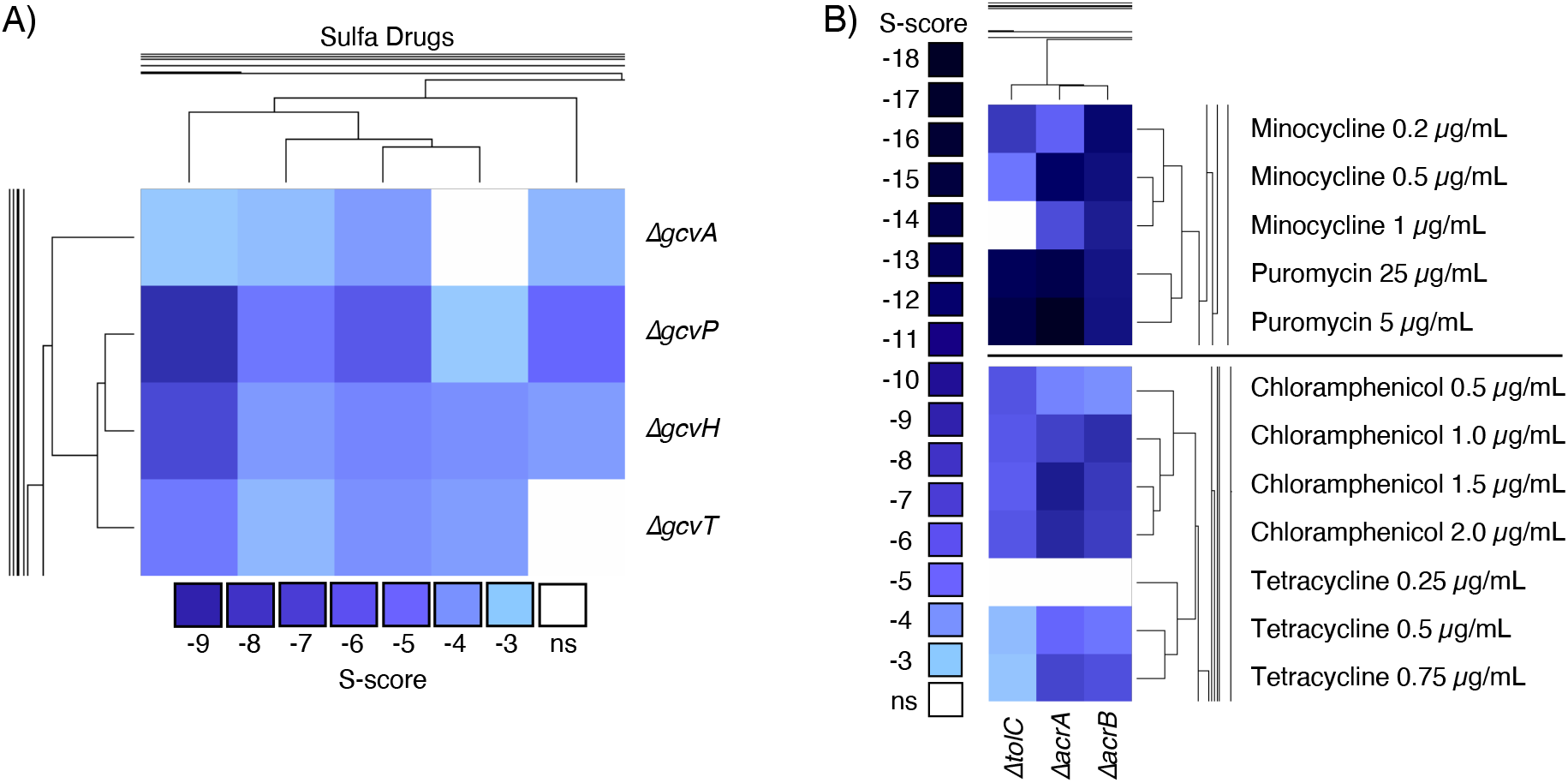
Clustered heatmaps displaying the S-scores for various single gene knockout mutants in different conditions. A) The GCV system in sulfonamide drugs (Sulfa), from left to right: Sulfamonomethoxine 100 μg/mL; Sulfamethoxazole 100 μg/mL; Sulfamethoxazole 200 μg/mL; Sulfamethoxazole 300 μg/mL; Sulfamonomethoxine 50 μg/mL. B) AcrAB-TolC system mutants in presence of AcrAB-TolC substrates. ns = non-significant.

Another means of validating the accuracy of the S-scores is by searching for known synthetic lethal pairs. For example, efflux pumps are one of the major drug resistance mechanisms within bacteria. In *E. coli*, the major RND efflux pump system responsible for exporting many classes of antibiotics, including tetracyclines and chloramphenicol, is the AcrAB-TolC system ^23,24^. Thus, we can assume that if this pump is deleted, the cell will be sensitive to its substrates. Following S-score analysis by ChemGAPP Big, we confirmed that Δ*acrA*, Δ*acrB* and Δ*tolC* were in fact sensitive to the systems substrates, with significant negative S-scores between −3 and −18 for minocycline, puromycin, chloramphenicol, and tetracycline (Fig. 5B).

The significant S-scores for both the GCV system and the AcrAB-TolC system further provide evidence that ChemGAPP Big is capable of producing accurate S-scores for successful hit acquisition.

### ChemGAPP Small produces informative fitness ratios

To validate ChemGAPP Small a screen was performed based on the observation within the Nichols KEIO chemical genomics dataset that Δ*envC* showed decreased fitness in membrane perturbing stresses ^4^. Within the Nichols dataset, Δ*envC* within 0.5% SDS + 0.5 mM EDTA had an S-score of −8.5, showing a significant decrease in fitness. In the current study, Δ*envC* was grown on LB and LB + 0.25% SDS + 0.25 mM EDTA. Within both conditions, a statistically significant decrease of Δ*envC* fitness was seen compared to the wildtype (Fig. 6). However, this decrease was more severe in the 0.25% SDS + 0.25 mM EDTA condition, with a fitness ratio of 0.28 compared to 0.77 on LB. This indicated a similar response to that observed in the Nichols et al. chemical genomic screen. Demonstration of this response by ChemGAPP Small, provides evidence for its effectiveness at interpreting small-scale screen data into informative fitness ratios.

**Figure 6:**
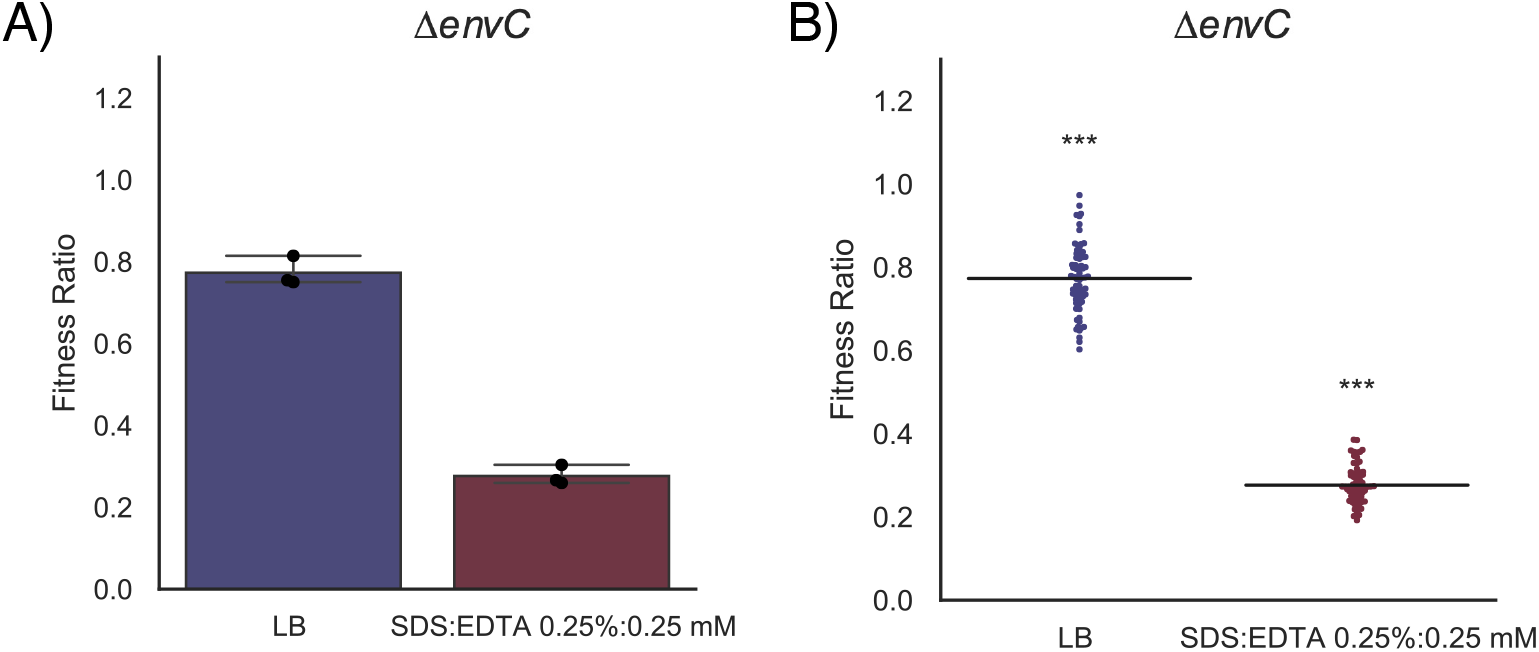
ChemGAPP Small produces informative fitness ratios. ChemGAPP Small provides the option of two different output plot types A) Bar plot output of ChemGAPP Small for Δ*envC* in LB, and 0.25% SDS + 0.25 mM EDTA, error bars represent 95% confidence intervals B) Swarm plot output of ChemGAPP small for Δ*envC* in LB, and 0.25% SDS + 0.25 mM EDTA. ns = p > 0.05; *** = 0.0001 < p ≤ 0.001.

### ChemGAPP GI successfully reveals genetic interaction types

To validate ChemGAPP GI we reanalysed data from a previous study evaluating the genetic interactions between the outer membrane (OM) lipoprotein *nlpI* and peptidoglycan machineries ^25^. Banzhaf et al., showed that *nlpI* did not genetically interact with the penicillin binding protein 4 (PBP4) encoding gene *dacB*. However, *nlpI* showed a synergistic interaction with PBP1B gene *mrcB* ^25^. ChemGAPP GI successfully reproduced these results using the raw data from the Banzhaf et al. study (Fig. 7A, 7B). In another study, a positive genetic interaction between the OM lipoprotein *bamB* and DNA replication activator *diaA* was described (Bryant et al., 2022). Using ChemGAPP GI the same conclusions were achieved (Fig. 7C). ChemGAPP GI, is consequently capable of accurately predicting all types of genetic interactions.

**Figure 7:**
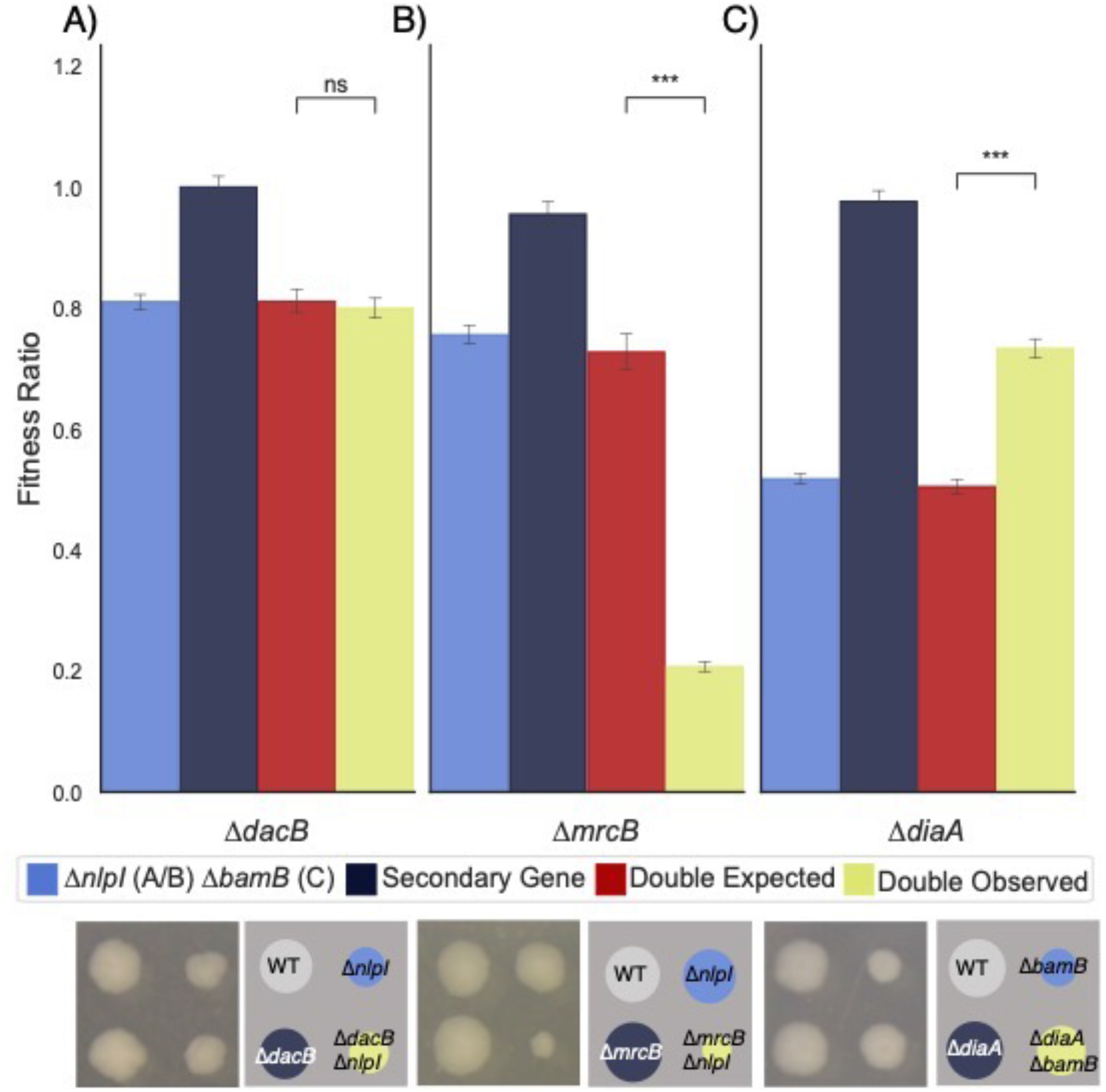
ChemGAPP GI can accurately predict all types of genetic interactions. A) Non-significant difference between the double expected and double observed fitness ratios, therefore indicating no epistasis between *dacB* and *nlpI*. B) Double observed significantly lower than double expected, showing negative epistasis between *mrcB* and *nlpI*. C) Double observed significantly fitter than double expected. Showing positive epistasis between *diaA* and *bamB*. ns = p > 0.05; *** = 0.0001 < p ≤ 0.001.

## Discussion

ChemGAPP was designed to address the need for a chemical genomics analysis software that is easy to use and fully suited to purpose. The current lack of software has led to the implementation of in-house scripts, often based on EMAP Toolbox ^16,18^. This gap in the field limits the analysis pipeline to bioinformaticians and computational biologists, who are able to write and implement these scripts. One recent study, however, produced an analysis package (ScreenGarden) for plate-based high-throughput screens to fill the gap of deprecated analysis software ^26^. However, similar to EMAP Toolbox, this software was not specifically designed for chemical genomic screen data and was tested on Synthetic Physical Interaction (SPI) screen data ^26^. Furthermore, ScreenGarden produces log growth ratios (LGRs) and Z-scores which are compared to one or two control plates as its fitness measure. For the analysis of chemical genomic screen data this is not as effective as using S-scores, which consider the effect of gene mutation across all conditions. By considering all condition effects, a more robust baseline to elucidate the true phenotype of a mutant within specific conditions is produced. Despite this, in instances where fitness ratios are more applicable, such as small-scale screens, ChemGAPP Small covers this requirement. This makes ChemGAPP a more comprehensive tool and, considering the nature and aim of chemical genomic screens, more suitable for chemical genomic analysis.

ChemGAPP is available in two formats at https://github.com/HannahMDoherty/ChemGAPP. Firstly, the software can be run through a Streamlit GUI, meaning no knowledge of coding or programming is required for end users. Secondly, the package is available as modules which can be run in command line, giving the user flexibility to choose or modify the features they wish to implement.

## Conclusions

We have introduced ChemGAPP a comprehensive, user-friendly wrapper software dedicated to the analysis of a variety of chemical genomic screen types. ChemGAPP’s three sub-packages were specifically designed to allow a wider scientific audience to engage in chemical genomic techniques by streamlining each dedicated analysis pathway.

## Methods

### ChemGAPP Big Pipeline

#### ChemGAPP Big: Conversion of Iris files

The package takes input files in the Iris format, which is a tab delimited tabular data, with columns specifying mutants’ coordinates on the plate and their growth features. Iris files were compiled into a dataset comprising of each condition replicate plate as the columns, and colony size data as the values. To initially improve the dataset, zero values where not all replicates were also zero were removed, since these likely represent mis-pinned colonies.

#### ChemGAPP Big: Plate Normalisation

Plate normalisation was performed as in Collins et al., 2006, with the addition of an initial check for edge effects to evaluate if the first normalisation step was required ^17^. Here, a Wilcoxon rank sum test was performed between the colony size distributions for the outer two edges and the centre of the plate. We opted for this test as the assumption of normality did not hold for the distribution of measurements. If edge effects are present, the distributions of colony sizes should differ. Only when distributions differ is the first stage of normalisation performed. In the second stage, all plates were scaled such that the plate middle mean was equal to the median colony size of all mutants within the dataset.

#### ChemGAPP Big: Quality Control Analyses

##### Quality Control Analyses: Z-score Test

The first test, aimed at distinguishing between the phenotype of no growth, and artefacts due to mis-pinned colonies or missed detection by IRIS is the Z-score test analysis. The Z-score value describes the distance of a test value from the population mean in measures of standard deviation. This adjusts the distribution of the population to a standard normal distribution, with a mean of zero and a standard deviation of 1 ^27^. Here, the Z-score test compares replicate colonies for each plate in each condition and highlights colonies which are outliers (Z-score value >1 or <-1) or missing values. ChemGAPP Big then calculates percentage normality for each replicate plate, i.e., the percentage of colonies within the plate which are not outliers or missing values.

##### Quality Control Analyses: Mann-Whitney Test

The next test, i.e., the Mann-Whitney test, assesses differences at the plate level. Here the distribution of colony sizes across each replicate plate are compared. If two replicates are reproducible, they should fall into the same distribution. This test is particularly useful for identification of mislabelled plates since distributions of colonies will be highlighted as significantly different. Furthermore, like the Z-Score test, the Mann-Whitney test can highlight unequal pinning effects. The Mann-Whitney test was performed on a plate-by-plate basis. For each replicate in the condition of interest, the distribution of mutant colony sizes was compared to each other replicates distribution and a Mann-Whitney *p*-value calculated. The mean *p*-value for each replicate was calculated by averaging the Mann-Whitney *p*-values which included the replicate of interest.

##### Quality Control Analyses: Condition Level Tests

Where all replicate plates are non-reproducible, a condition can be deemed unsuitable for screening. This can occur, for example, when a condition produces a coloured media which is difficult to distinguish from the colonies. In such cases Iris can struggle to differentiate the colonies from the background media, leading to artefacts in detection ^3^. Therefore, it is important to also consider if a condition as a whole should be removed from the dataset. Here two tests were implemented, the first calculated the variance in the Mann-Whitney p-values for each replicate plate. A large variance here suggests that the entire condition may be detrimental, since it is likely all replicate distributions differ from each other. Finally, a general variance analysis was employed. Here, the variance of each mutant replicate in the condition of interest was calculated. The mean of these variances was then calculated to gain an average variance across all plates in each condition. A high variance implies that most colony replicates differ within their size, and thus the entire condition is non-reproducible.

##### ChemGAPP Big: Threshold selection and data removal

For each of the plate level statistical analyses, the percentage of plates that would be lost within a condition at certain thresholds was calculated. For Z-score analysis data, thresholds of 20%, 30%, 40%, 50%, 60%, 70% and 80% for normal colonies were tested. For the Mann-Whitney test results thresholds were generated based on the obtained range of mean p-values. For the condition level analyses, the number of conditions that would be lost at different thresholds was calculated. The thresholds chosen were generated based on the obtained range of variance values for both condition level tests. The chosen thresholds for the reanalysed KEIO dataset were 80%, 0.07585, and 0.0128, for normality, Mann-Whitney plate analysis and Mann-Whitney condition analysis. Data was not removed based on the average variance test since little variance within conditions was observed (Fig. S1). Chosen thresholds were inputted and a new dataset with failing plates and conditions removed was outputted and normalised.

##### ChemGAPP Big: Fitness Score Calculation

Colony size data was converted to fitness scores (S-scores) via a modified t-value score, as described in Collins et al., 2006, this was: ^17^

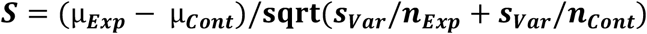

Where: μ_Exp_ = the mean of normalised colony sizes for the mutant of interest in the condition of interest; μ_Cont_ = the median of normalised colony sizes for the mutant of interest in all conditions; n_Exp_ = number of replicates of mutant of interest in condition of interest; n_Cont_ = median number of replicates for all conditions. S_Var_ is equal to:

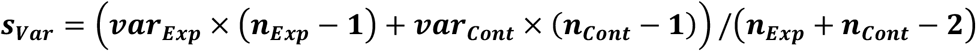

Where: var_Cont_ = median variance of normalised colony sizes for mutant of interest in all conditions or a minimum bound; var_Exp_ = the variance of normalised colony sizes for mutant of interest within the condition of interest or a minimum bound. The minimum bound for var_Exp_ was placed on the expected standard deviation used to calculate var_Exp_. The average standard deviation for all mutants within the condition of interest, was used to produce a minimum bound variance for var_Exp_.

The minimum bound for var_Cont_ was equal to μ_Cont_ multiplied by the median relative error of all mutants in all conditions, where:

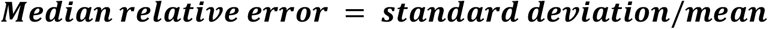

Where S-scores were calculated as infinity, scores were converted to missing values. The S-scores of each plate were then scaled such that the interquartile range (IQR) of the scores was equal to 1.35, matching that of a standard normal distribution. This scales the datasets such that the values for significant hits are for S-scores < −3 and > 2. In general, the majority of S-scores within a dataset will be close to zero. Significant hits are S-scores that are classified as outliers. Outliers are commonly determined as scores deviating from the first or third quartiles by 1.5 times the IQR, which in this case is ~2. Therefore, significant scores would be classified as < −2 and >2. However, since loss of function mutants are more common, a higher threshold of deviation was chosen to represent significant negative S-scores. The final non-curated dataset was visualised within Treeview3 ^28^. Data was hierarchically clustered for both rows and columns using uncentered Pearson Correlation and average linkage.

##### ChemGAPP Big: Benchmarking the curated dataset

The non-curated and curated datasets were compared to each other as well as to the Nichols et al., 2011 KEIO dataset, which was analysed within EMAP Toolbox ^4^. Genes from the same operon and genes from differing operons were subjected to cosine similarity analysis. Phenotypic profiles that are similar will have cosine similarity scores close to 1, those with no similarity will have scores close to zero, and negatively correlated profiles will have scores close to −1. Confusion matrices were produced at different thresholds of cosine similarity score, ranging from −1 to 1, by increments of 0.1. Here, a true positive was a set of genes from the same operon with a cosine similarity score greater than the threshold; a true negative was a set of genes from different operons with a cosine similarity score lower than the threshold; a false positive was a set of genes from different operons with a cosine similarity score greater than the threshold; and a false negative was a set of genes from the same operon with a cosine similarity score lower than the threshold. At each threshold the sensitivity, specificity and false positive rate were calculated and plotted in an ROC curve. The AUC for the datasets was then calculated and compared.

##### ChemGAPP Big: Production of Bootstrapped Dataset

In order to evaluate the accuracy and robustness of the S-scores produced by ChemGAPP Big, a bootstrapped dataset was produced. The KEIO non-curated and curated normalised datasets were bootstrapped 1000 times with replacement. Each of the 1000 datasets was subjected to S-score analysis. The mean of the bootstrapped S-scores for each mutant in each condition, were compared to the true S-scores and the mean absolute error (MAE) calculated.

### ChemGAPP Small Pipeline

The small-scale Iris files were compiled and normalised as for ChemGAPP Big; however, zero values were not converted to missing values. Fitness ratios were computed for each plate either by dividing the mean mutant colony size by the mean wildtype colony size for the bar plots, or by dividing each individual colony by the mean wildtype colony size for the swarm plots. Significance for bar plots was measured by 95% confidence interval, whereas a one-way ANOVA and Tukey HSD analysis was used to determined significant differences from the wildtype within the swarm plots.

### ChemGAPP GI Pipeline

Genetic interaction Iris files from Banzhaf et al. 2020 and Bryant et al. 2022, were inputted into ChemGAPP GI and genetic interaction fitness ratios calculated. For the single and double knockouts, genetic interaction fitness ratios were calculated by dividing the colony size of the mutant by the colony size of the WT for that replicate. The double knockout expected fitness ratio was calculated by the formula:

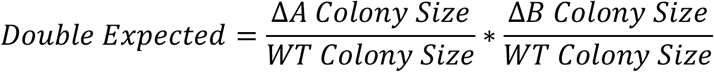

Significance was measured by 95% confidence interval error bars, and one-way ANOVAs with Tukey HSD analysis between the double observed and double expected fitness ratios.

## Supporting information

Dependencies for software

## Abbreviations

ROC: Receiver operating characteristic curve
AUC: Area under the curve
MAE: Mean absolute error
STD: Standard deviation
THF: Tetrahydrofolate
OM: Outer membrane
IQR: interquartile range
Sulfa: Sulfonamide drugs

## Declarations

### Ethics approval and consent to participate

Not applicable.

### Consent for publication

Not applicable.

### Availability of data and materials

All data generated or analysed during this study is included in this published article and its supplementary information files.

The benchmarked dataset produced by Nichols et al is available within the supplementary information files of:

Nichols, R. J. *et al*. Phenotypic landscape of a bacterial cell. *Cell* **144**, 143–156 (2011). DOI: 10.1016/j.cell.2010.11.052

Project name: ChemGAPP

Project homepage: https://github.com/HannahMDoherty/ChemGAPP

Operating system: Platform independent

Programming language: Python

License: by-nc-nd-4.0

Requirements:

- Python3
- Other dependencies listed within Additional File 3, found within the supplementary information.

Source code for ChemGAPP is available on GitHub at https://github.com/HannahMDoherty/ChemGAPP

### Authors’ contributions

HMD, MB, and DM conceived and designed the study. HMD wrote the manuscript, developed the software, and ran all data analyses. MB performed the *envC* chemical genomics screen. GK and MG supported the writing of the code. MB, DM, and MG helped edit the manuscript. All authors reviewed and approved the manuscript.

## Acknowledgements

The authors thank Athanasios Typas for assisting the project.

## Supplementary information

Additional File 1 (pdf). Supplementary Figures. Figures S1 and S2.

Additional File 2 (pdf). Supplementary Methods. Extended methods for plate normalisation, production of the bootstrapped dataset, and the small-scale envelope stress screen.

Additional File 3 (txt). Dependencies for ChemGAPP.

