## Supplementary Figures for "ChemGAPP; A Package for Chemical Genomics Analysis and Phenotypic Profiling"

### ChemGAPP Big

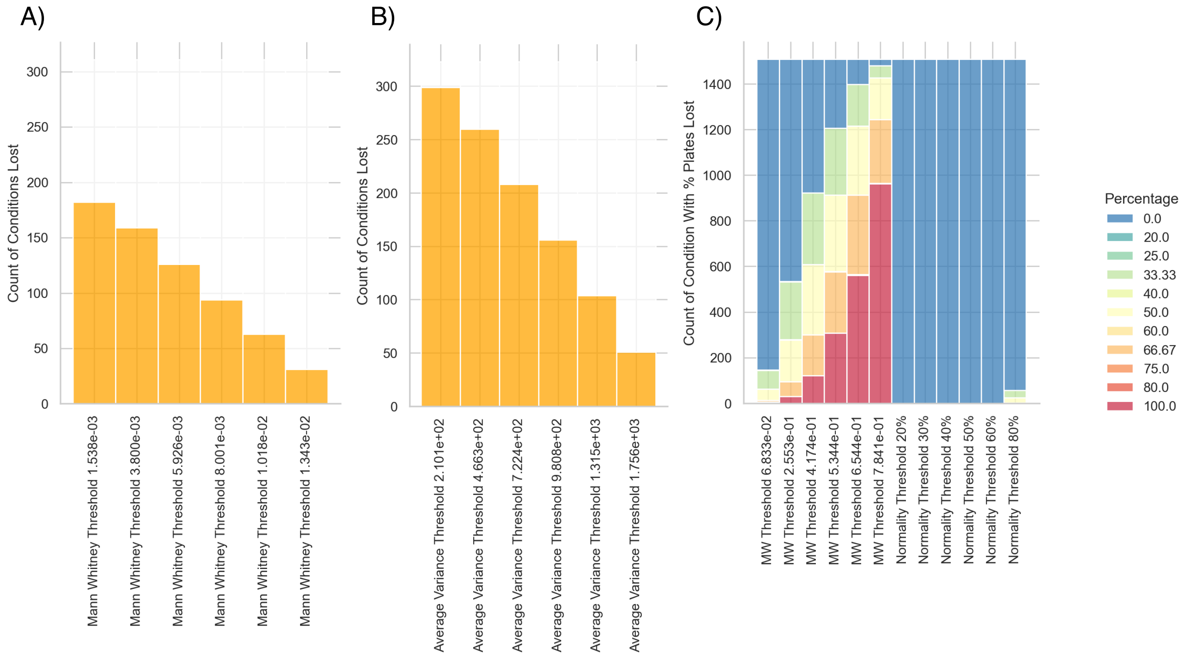


**Figure S1: Threshold selection bar plots outputted by ChemGAPP Big**. ChemGAPP Big makes threshold selection simple, with informative plots which display the quantity of data lost at various threshold for: A) Mann-Whitney Condition Level Analysis; B) Condition Level Variance Analysis; C) Mann-Whitney Plate Level Analysis and Z-score analysis.

**ChemGAPP Big identifies common errors within chemical genomic screens.**

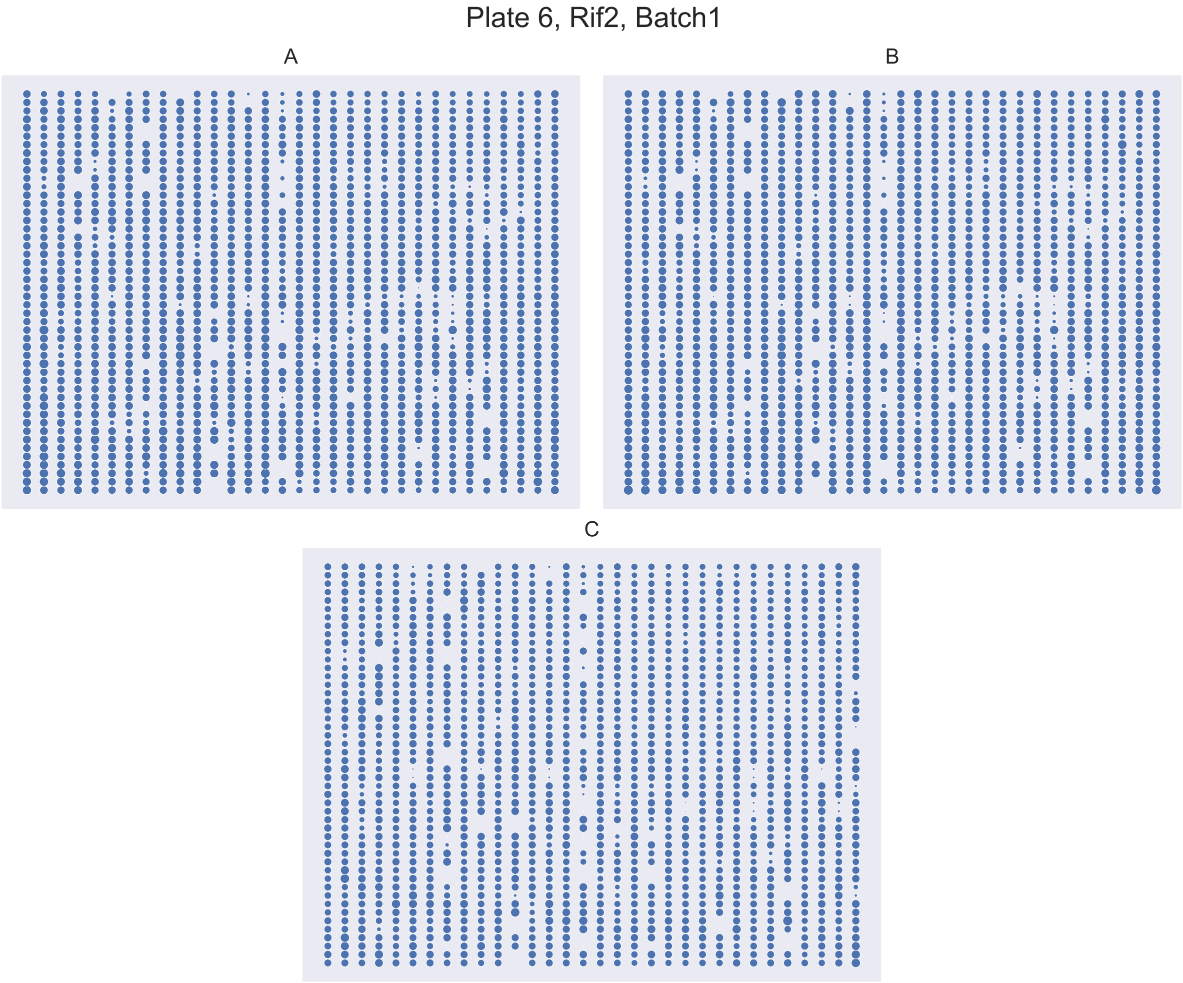


**Figure S2: Replicate C highlights defects of missed and unequal pinning**. Plate matrix depicting the colony sizes within replicate plates of the condition Rif2 (Rifampicin 2 µg/mL), Plate 6, Batch1. Replicate C shows an increased number of missing colonies (21) compared to A (4) and B (4). The upper segment of replicate C has generally smaller colonies than the lower segment, highlighting unequal pinning.
