## Supplementary Methods for "ChemGAPP; A Package for Chemical Genomics Analysis and Phenotypic Profiling"

**Plate Normalisation**

Plate normalisation was performed for each plate in two stages as in Collins et al., 2006 [1]. The outer two edges of the plate were first checked for to see if normalisation was required to remove edge effects. The Wilcoxon rank sum test was performed between the outer two edges and the centre of the plate. Where $p < 0.05$ distributions were considered as different, and the normalisation was performed. Here, medians for each edge row and column were calculated and colony sizes scaled such that these medians were equal to the plate middle mean (PMM), as such:

$$\boldsymbol{Normalised\ size} \\ = \boldsymbol{Unnormalised\ size} \times (\boldsymbol{PMM/Median\ of\ row\ or\ column})$$

The PMM corresponds to the mean of colonies, excluding the outermost rows and columns, that fall within the 40$^{th}$ and 60$^{th}$ percentiles of colony size. In the second stage all colonies were normalised. The median colony size for all colonies was calculated and the PMM scaled to equal this median value, as such:

$$\boldsymbol{Normalised\ size} = \boldsymbol{Unnormalised\ size} \times (\boldsymbol{Plate\ median/PMM})$$

**Production of Bootstrapped Dataset**

The normalised curated and non-curated KEIO datasets were bootstrapped 1000 times, each time 300 conditions were randomly selected with replacement. S-scores were calculated, without gene assignment, for each bootstrap and the average s-score across all bootstraps was calculated for each mutant in each condition. Due to the bootstraps being produced with replacement, on some occasions all replicates of specific conditions were identical. In these instances, $var_{Exp}$ was equal to zero and as such the minimum bound also zero. Further to this, for a small number of genes $\mu_{cont}$ was also equal to zero, since these mutants were synthetically lethal in most conditions. Due to these three zero values, a small number of abnormally large S-scores were produced. Since this is an artifact of the bootstrapping method and would not occur within the normal confines of ChemGAPP Big, bootstrapped S-scores > 125 were removed as outliers. Exclusion was performed prior to S-score averaging. Excluded values represented a negligible 0.000018%, and 0.000027% of all values within the bootstrapped non-curated dataset, and the bootstrapped curated dataset, respectively. S-scores > 125 within the original curated dataset were also removed, for ease of visualisation. However, these scores represented just 0.00056% of all values within the dataset. The absolute error was then computed between the original dataset s-scores and the average bootstrapped s-scores for each mutant in each condition, for both the curated and non-curated datasets. The mean absolute error (MAE) was then calculated for both datasets. The calculation of the absolute errors for the curated dataset included original curated S-scores > 125, but not Bootstrapped S-scores >125.

**ChemGAPP Small: Envelope stress screen**

Wildtype BW25113 and BW25113 Δ*envC* were pinned onto LB agar, and LB agar supplemented with 0.25% SDS + 0.25 mM EDTA. Three replicates for each plate were produced and grown for 12 hours at 37˚C before imaging. Plate images were analysed using the image analysis software IRIS and the colony size phenotype was measured [2].

1. Collins, S. R., Schuldiner, M., Krogan, N. J. & Weissman, J. S. A strategy for extracting and analyzing large-scale quantitative epistatic interaction data. *Genome Biology* **7**, R63 (2006).

2. Kritikos, G. *et al.* A tool named Iris for versatile high-throughput phenotyping in microorganisms. *Nat Microbiol* **2**, 17014 (2017).
